# 3D Printed Bioelectronic Scaffolds for Impedance-based Cytotoxicity Monitoring of In Vitro Cancer Models

**DOI:** 10.64898/2026.05.07.719019

**Authors:** Somtochukwu S. Okafor, Sandra K. Montgomery, Jae Park, Tianran Liu, Mbama Safrega, Justin S. Yu, Cayleigh P. O’Hare, Angela Schab, Anna P. Goestenkors, Cielo J. Vargas Espinoza, Yuqing Wu, Elena Lomonosova, Ismael Seáñez, Mary M. Mullen, Alexandra L. Rutz

## Abstract

Cancer is a significant contributor to global mortality and places a substantial burden on healthcare systems, underscoring the need for improved strategies for developing and evaluating new therapies. Electrochemical impedance monitoring of in vitro cancer models is a promising technique for evaluating treatment effectiveness, particularly for evaluating how well a drug may kill cancer cells. This approach is advantageous over conventional end-point assays because it is non-destructive, label-free, and can provide temporal information on cell behavior and drug kinetics. However, traditional impedance devices are limited in that they do not support three-dimensional cell culture that has become standard in cancer studies. Typical devices are planar substrates that support monolayer culture, which has been shown to overestimate drug effectiveness. In this work, we propose 3D printed bioelectronic scaffold devices that provide 3D cancer cell culture while functioning as an on-chip readout for monitoring changes in cell characteristics via impedance. We describe device development and demonstrate reproducible fabrication, stable electrochemical properties, cell detection by impedance, and proof-of-concept monitoring of cytotoxicity in response to a chemotherapeutic drug. Overall, this technology offers a promising platform that could be further developed for compound screening as part of drug development or precision medicine.

## Introduction

Cancer is a major cause of death worldwide and was responsible for 9.7 million deaths globally in 2022^1^. Significant advancements in cancer research have led to the development of novel drug compounds targeting several cancer types. However, despite this progress, there is still a critical need to develop additional novel therapeutics, owing to the complexity of many cancers^2^. Furthermore, tumor heterogeneity and patient-to-patient variability can lead to poorly effective treatments when these are broadly applied to an entire population of a given cancer type^3^. Precision medicine can solve this challenge by tailoring therapy to an individual after identifying compounds that result in maximal drug effectiveness using their own cells^3–6^. Both of these areas – drug development and precision medicine – require cancer models for drug screening. In vitro models of cancer are favored for ease of generation, eliminating use of animals, and human-specific biology^7,8^. In vitro cancer models can range from simple monolayer culture on tissue culture plastic to 3D culture models such as spheroids or organoids to more sophisticated systems relying on hydrogels, scaffolds, and/or tissue-on-a-chip devices^3,9–12^. For cancer especially, three-dimensional culture is strongly favored over 2D monolayer culture for several reasons. In the context of drug testing, 2D culture models are unable to recapitulate the properties of in vivo tumors including their resistance to therapies^5,13–15^, often leading to over-estimation of therapy effectiveness and reduced translatability of studies^16,17^.

It is anticipated that reliance on in vitro models will continue to expand in commercial and clinical settings. As one example promoting this shift, the United States Food and Drug Administration Modernization Act 2.0 removed animal model mandates for drug development and now accepts in vitro models as an alternative^13^. With the anticipated rise of these models, quantitative and high throughput methods of measuring biological read-outs are needed^18,19^. For anti-cancer drug screening specifically, a read-out of drug response is needed and that is most commonly cytotoxicity of the cancer cells^4,20^. Many cytotoxicity assays have been developed and commercialized, and these include metabolic activity assays such as 3-(4,5-dimethylthiazol-2-yl)-2,5-diphenyltetrazolium bromide (MTT), water-soluble tetrazolium salt (WST), and adenosine triphosphate (ATP)-based assays^20^ as well as cell membrane integrity assays such as live/dead staining.

While such approaches generally provide straightforward readouts with developed protocols, notable limitations exist. Cytotoxicity assays are generally developed for use on 2D cell culture and even when adapted for 3D cultures preferred for cancer, they face several shortcomings. Optical (fluorescence, absorbance or luminescence) – based assays can be limited by poor dye penetration into dense 3D cultures^21^. Imaging-based assays may additionally suffer from insufficient sampling due to limited imaging depth or may require time-consuming and technically demanding protocols in order to more holistically image the sample^21^. Additionally, these assays are typically performed as an end-point analysis; however, temporal information could be valuable. Different drugs, or different doses of the same drug, can vary in their kinetics, with some inducing decreased cell viability (cytotoxic or cytostatic effect) more rapidly than others depending on their mechanism of action^20^. Some cells may initially die in response to treatment, but other surviving cells may later develop resistance^22^, mimicking the therapeutic resistance seen in patients. Studies have also indicated that drug efficacy may increase after a washout period (removal of drug)^23^. This disregarded information may be valuable in decision-making for precision medicine or may offer insights into mechanisms of action in novel compound screening.

Cell-based electrochemical impedance monitoring could serve as an alternative assay for evaluating cytotoxicity of cancer models^24–27^. Impedance is a frequency dependent measure of the resistance to flow of alternating current. In impedance monitoring utilizing a two-electrode configuration, a small sinusoidal potential is applied between a working electrode and counter electrode over a range of frequencies (typically ∼10^-1^-10^5^ Hz). The resulting current is measured and the impedance is calculated using Ohm’s law^28^. When cells are seeded and adhere onto the working electrode surface, the insulating properties of their membranes act as a barrier, thereby impeding current at the electrode-solution (cell culture media) interface. Thus, the electrode impedance increases with increasing cell coverage^24,29–31^. In addition to changes due to cell coverage on the electrode, information on cell-cell contacts such as integrity of tight junctions, changes in cell morphology, and cell-electrode interactions including adhesion strength can be extracted from the impedance at different frequencies^24,28^. In all, electrical impedance serves as a valuable and versatile tool for assessing cell behavior.

Several studies have employed impedance monitoring to investigate the cytotoxic effects of chemotherapeutic drugs on cancer cells, demonstrating accuracy comparable to established methods^24,25,32–34^. Notably, some studies have reported higher sensitivity with impedance-based detection compared to other techniques^35^. Impedance monitoring involves using electrodes as not only the device for providing the on-chip readout of cell characteristics but also as the culture platform since the measurement depends on cell-electrode interfacing. In the majority of these works, cells are grown on planar electrode arrays patterned on a glass, silicon, or plastic substrates^24,28^. Such surfaces provide a 2D culture substrate that is not ideal for culturing cancer models, as discussed above^24,36–38^. Alternatively, electrodes can be designed to simultaneously support 3D culture, while providing the desired electronic impedance readout. 2D plates of electrode arrays have been altered to enable spheroid or organoid cultures and vertical electrodes have been employed for 3D impedance sensing^39–44^. These systems have been shown to detect cytotoxicity in response to the addition of chemotherapeutic drugs for several cancers^39,42–44^. However, a limitation with this approach is lack of direct contact between the majority of the cell volume and electrode^21,36^. Another approach is to construct the 3D culture substrate from an electronically conducting material. Scaffolds built from conducting polymers have been developed into 3D devices for cell-based impedance monitoring^45–48^, primarily for evaluation of gut barrier function. Yet, the potential of conducting 3D scaffolds for monitoring cytotoxicity of cancer cells over time after administration of drug compounds remains largely unexplored.

Building off our previous work of developing bioelectronic scaffolds – 3D printed conducting hydrogel scaffolds^49^, we envisioned using these structures directly as electrodes for 3D cell culture impedance monitoring by mounting the scaffolds onto electrical connections. In this work, we investigate if bioelectronic scaffolds can support 3D cancer cell culture and when used as an electrode, can detect the presence of cells through impedance measurements (Figure 1). We also investigate if the bioelectronic scaffold electrodes can detect changes in cell number over time corresponding to cell proliferation and chemotherapy drug-induced cell death. Overall, our results highlight the potential of these bioelectronic scaffolds to be used as 3D impedance sensors for monitoring cytotoxicity over time, which could be useful in the future for drug screening as part of precision medicine or drug development.

**Figure 1:**
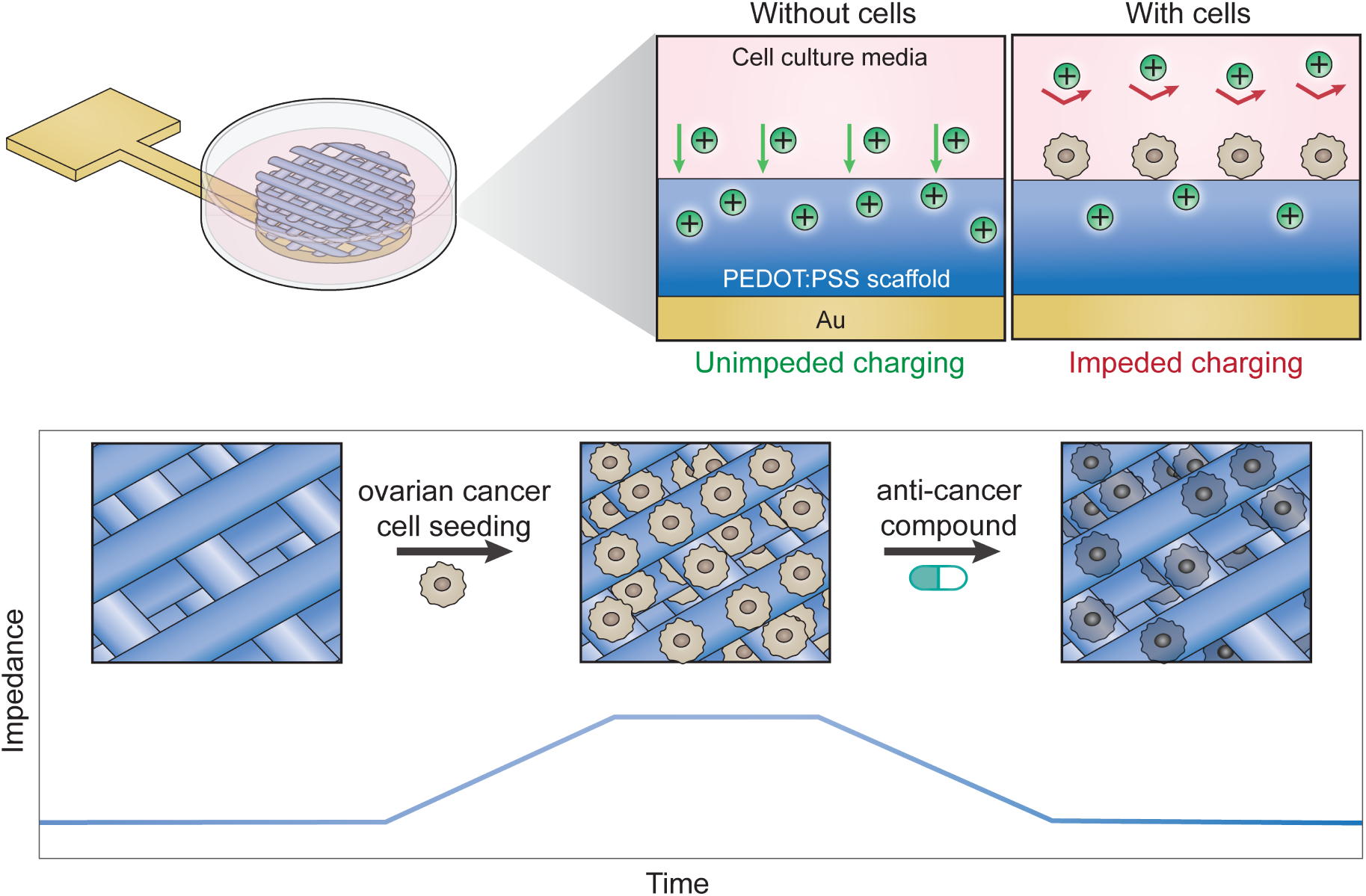
Overview of 3D printed bioelectronic scaffold electrodes for impedance-based cytotoxicity monitoring of in vitro cancer models. When cells are seeded and attached to bioelectronic scaffold electrodes, they act as a barrier and impede current flow at the electrode/media interface. As cells proliferate, the impedance of the electrode increases further, relative to its cell-free state. When an anti-cancer drug compound is introduced, this results in cell death. As cancer cells die, they detach from the scaffold or degrade causing a decrease in impedance approaching its cell-free values.

### Bioelectronic scaffold electrodes have stable electrochemical properties in cell culture conditions

Impedance monitoring of cells typically employs a two-electrode configuration consisting of a working electrode and a counter electrode. The working electrode is where cells are seeded and is the site where electrochemical interactions are monitored to study cell characteristics (e.g. cell-cell interactions, cell-substrate interactions^28^). The counter electrode completes the circuit to apply a small alternating sinusoidal potential, in this case 10 mV across 10^-1^ – 10^5^ Hz frequencies. Here, we proposed our previously developed 3D conducting scaffolds^49^ as the working electrode. These conducting scaffolds were fabricated by 3D printing hydrogels of the conducting polymer poly(3,4-ethylenedioxythiophene) polystyrene sulfonate (PEDOT:PSS).

These scaffolds, however, required electrical connection to use as an electrode. We proposed bonding the scaffold to gold patterned on glass slides to provide a dry and stable electrical connection to the power supply (Figure 2A-B, Supplementary Figure 1). Inspired by the methodology of Savva et al.^45^, we utilized a thin film of PEDOT:PSS as an intermediary layer between the gold and PEDOT:PSS scaffold. This PEDOT:PSS adhesion layer contains 4-arm poly(ethylene glycol) thiol (PEG-SH) to provide a means of attachment of the film to gold (thiol-gold bonding). After bonding scaffolds, the scaffolds maintained their 3D and porous structure (thickness ∼500 μm) (Supplementary Figure 2). Additionally, after gentle agitation, scaffolds remained connected to the substrate and therefore appeared to be stable for cell culture processes like media changes. Opposite to these scaffold working electrodes, a large gold square (400 mm^2^ area) served as the counter electrode (Figure 2A, Supplementary Figure 1). Finally, a plastic chamber and lid were fixed around the electrodes to provide a closed system for aseptic in vitro study. Upon completion, each device comprised of 1 counter electrode and 3 working electrodes to provide biological replicates.

**Figure 2:**
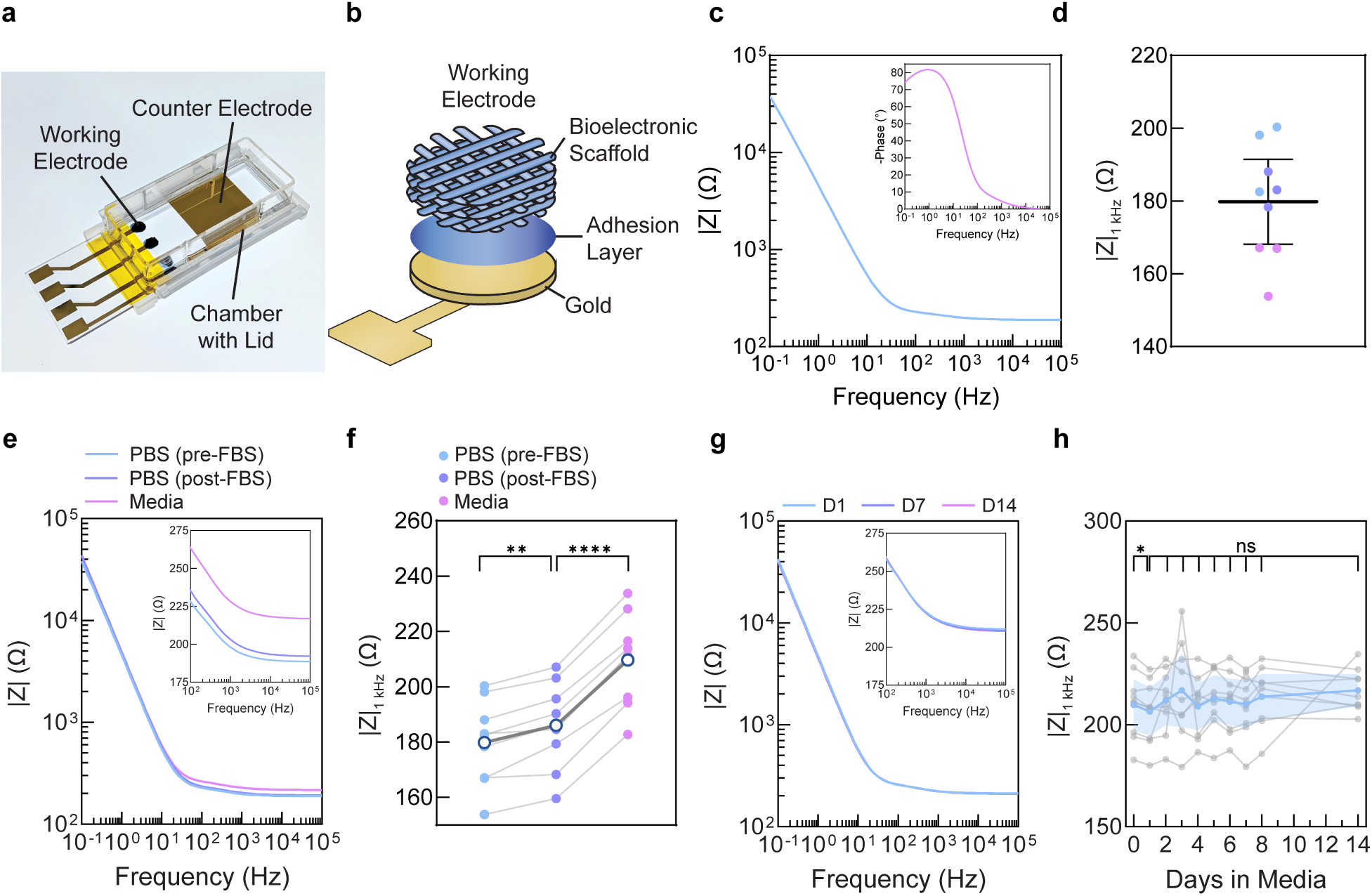
Bioelectronic scaffold electrodes have stable electrochemical properties in cell culture conditions. **a)** Photograph of assembled device. **b)** Schematic illustrating working electrode: 3D printed PEDOT:PSS scaffold bonded to gold. **c)** Representative Bode impedance magnitude (|Z|) plot and corresponding phase plot (inset) of the bioelectronic scaffold electrodes across multiple frequencies in phosphate buffered saline (PBS). **d)** Plot showing the distribution of the impedance magnitude at 1 kHz across all devices. Color of points indicates a device, each for which there are 3 electrodes (N=9 electrodes). Mean and 95% confidence interval presented. **e)** Representative Bode plots of |Z| in PBS (before and after fetal bovine serum (FBS) incubation) and in media (RPMI 1640 with 10% FBS). Inset shows zoomed in plots at 10^2^ – 10^5^ Hz. **f)** Quantification of changes in impedance magnitude at 1 kHz across all conditions identified in (e). N=9 electrodes. Mean depicted by white circles with dark blue borders and individual electrodes in color with repeated measures connected by a line. Repeated measures one-way analysis of variance (ANOVA) and Šidák-adjusted multiple comparison tests performed. ****P≤0.0001, ***P<0.001, **P≤0.01, *P≤0.05. **g)** Representative Bode plots of |Z| on 1, 7 and 14 days after incubation in cell culture conditions (RPMI 1640 + 10% FBS cell culture media, 37 °C, 95% relatively humidity, and 5% CO2). **h)** Impedance magnitude at 1 kHz in cell culture conditions over time. Individual electrodes presented in grey, mean (solid blue line) and 95% confidence intervals (blue shaded region) presented. N=9 electrodes. Repeated measure one-way analysis of variance (ANOVA) and Šidák-adjusted multiple comparison tests performed. *P≤0.05, non-significant (NS) P*>*0.05.

Following device fabrication, electrode properties were characterized in phosphate buffered saline (PBS) using electrochemical impedance spectroscopy. Working electrode impedance was measured to validate electrical connection to scaffolds, to confirm expected electrochemical properties of these PEDOT:PSS electrodes, and to determine baseline device properties prior to introduction of biology. The Bode impedance magnitude plot exhibited strong frequency dependence at low frequencies (<10 Hz), indicative of capacitive behavior, and a transition to more resistive (less frequency-dependent) behavior at higher frequencies (Figure 2C). This characteristic “hockey stick” profile is consistent with impedance spectra reported for similar electrodes^45,50,51^. Variability in electrode impedance within (intra) and across (inter) devices was then characterized to determine if the fabrication process is reproducible. Variability here is defined using the coefficient of variation (CV: standard deviation divided by the mean). In bioanalytical tools, a CV less than 10 - 15% is typically considered a good threshold for reproducibility of results^52^. In this study, 1 kHz, a frequency relevant for impedance-based cell monitoring^53^, was used for all device characterization unless stated otherwise. Among three devices, a CV of 5.0, 6.0, and 7.4% was observed, corresponding to the variation of impedance among each device’s three working electrodes. For all nine electrodes of the three devices combined, impedance values ranged from 153.8 to 200.3 Ω with a mean of 179.8 ± 15.19 Ω and the CV was 8.4% (Figure 2D). These results indicate reproducible manufacturing within and across devices for impedance monitoring.

The above characterization was performed on scaffold electrodes disinfected with 70% ethanol and washed with phosphate buffered saline solution (PBS). However, devices undergo additional preparation for cell culture; this includes (1) incubation in fetal bovine serum (FBS) to support cell attachment^49,54^, (2) PBS washing to remove excess serum, and (3) equilibration in the specific cell culture media (Supplementary Figure 3). We therefore sought to investigate how FBS and media incubation influenced electrochemical properties, as our previous work showed that PEDOT:PSS hydrogels adsorb media components^55^. Impedance magnitude significantly differed across the solution incubations as determined by a one-way repeated measures ANOVA (F = 514.1, p<0.0001). Pairwise comparison shows that after FBS incubation and return to PBS, impedance increased by 3.5 ± 2.0% relative to PBS before FBS (Figure 2E-F; p=0.0013, 95% CI [-9.347, -3.064]). Furthermore, the immediate change in measurement solution from PBS (after FBS incubation) to media (RPMI 1640 with 10% FBS) caused an additional 13 ± 1.9% increase in impedance when taken immediately upon solution change (Figure 2E-F; p<0.0001, 95% CI [-26.63, -20.82]).These findings collectively indicate that to confidently determine the baseline impedance prior to cell addition, impedance should only be measured after all preparative processes and additionally in the final cell culture media to be used throughout the study.

For monitoring cytotoxicity, it is important to determine if the scaffold electrode impedance remains stable or changes over time in cell culture conditions. Changes in impedance are the proxy measurement for cytotoxicity, and therefore, drift in impedance could be erroneously mistaken for cell changes if not corrected for. As a first step towards evaluating stability, the impedance of the bioelectronic scaffold electrodes did not change significantly in PBS for at least 56 days (Supplementary Figure 4; F = 0.7631, p = 0.7128, ordinary one-way ANOVA). To fully achieve impedance-based cell monitoring conditions, devices were incubated in RPMI 1640 with 10% FBS maintained in an incubator (37 °C, 5% CO_2_) and measured daily over two weeks. A one-way repeated measures ANOVA determined that the impedance magnitude did not significantly change over the two weeks (F = 1.060, p = 0.3778). Despite a non-significant effect of time, we still conducted pairwise comparisons given the fact that impedance monitoring of cells would be conducted day-to-day. A significant difference was observed from day 0 to day 1 (Figure 2H; p = 0.0429, 95% CI [0.2876, 6.084]). After that initial change, no further significant changes were observed between timepoints over the study duration (Figure 2H; p > 0.05). Thus, we identified that cell culture monitoring could be started as soon as one day after devices were introduced to and equilibrated with cell culture conditions. After one day, there was no evidence of drift in the impedance over time; a linear regression yielded a slope not significantly different from zero (Supplementary Figure 5D; slope = 0.3155, R^2^=0.006042, F = 0.4802, p = 0.4903). There was low day-to-day variability in the mean impedance of nine electrodes; the coefficient of variation was 2% (Supplementary Figure 5A-C). Thus, given the absence of any time-dependent changes in mean impedance, this day-to-day variability was interpreted as random fluctuations, herein referred to as noise. In summary, these various analyses give us confidence to move forward with impedance monitoring of cells after thoroughly characterizing baseline device properties as a function of preparation, solution type, and time.

### Identifying optimal frequency for impedance-based detection of cells on bioelectronic scaffold electrodes

In our prior work, we demonstrated that these 3D printed PEDOT:PSS scaffolds possessing an interconnected porosity supported high cell viability and proliferation of human dermal fibroblasts^49^. Towards our goal of impedance-based cytotoxicity monitoring of cancer models, we first sought to establish that these scaffolds could support a different cell type - ovarian cancer cells. OVCAR8, a human ovarian cancer cell line derived from high grade serous ovarian carcinoma, was selected as the model as it is widely used to investigate mechanisms of drug resistance as well as to screen and develop novel therapeutics^56,57^. To promote a dense cell culture mimetic of tumors, we leveraged the customization afforded by additive manufacturing to design the scaffolds with small pores to promote pore filling and dense cell packing. The approximate dimensions were as follows: diameter 4 mm, pore size 90 μm, thickness 720 μm (6 layers; Figure 3A).

**Figure 3:**
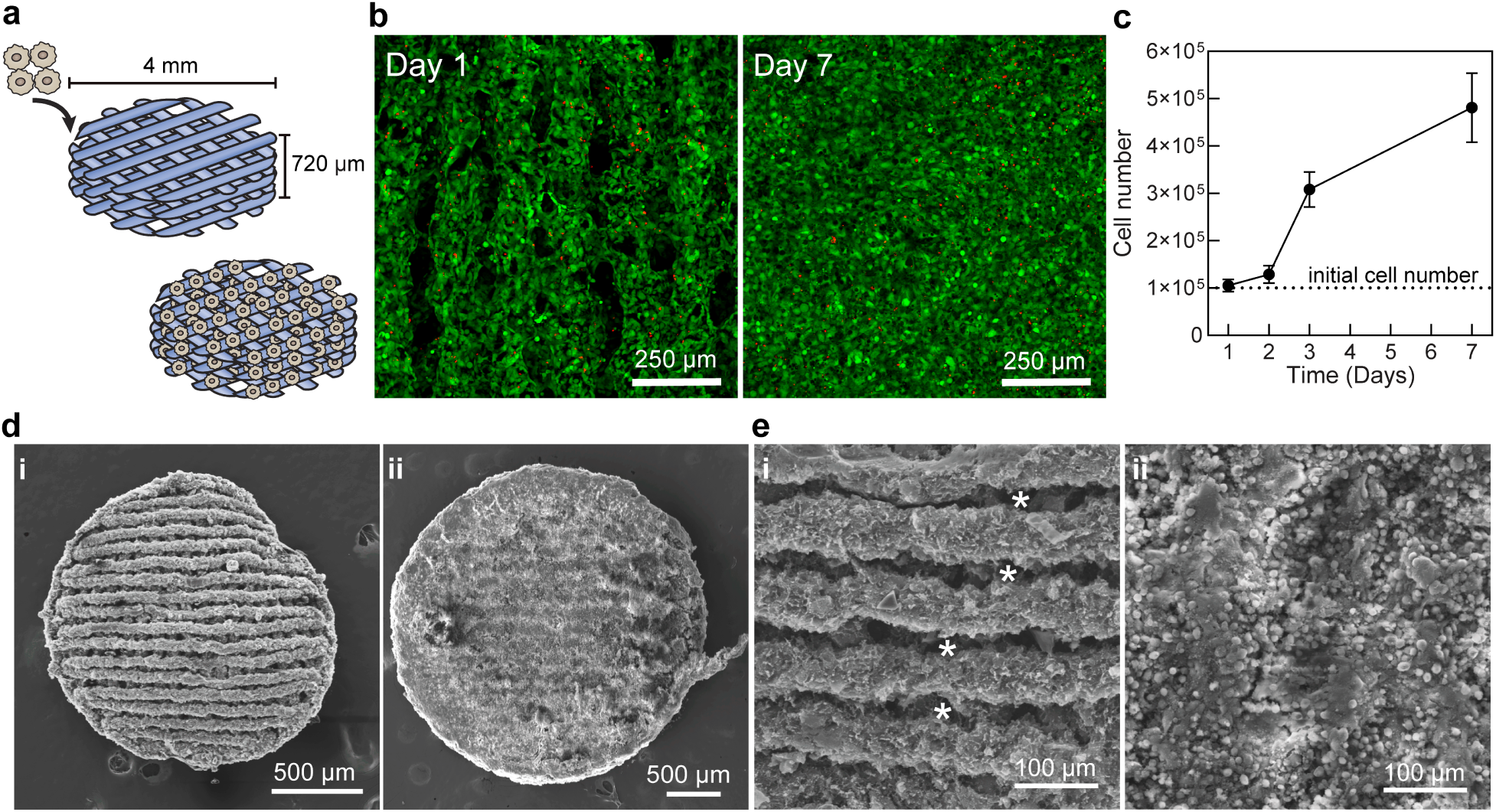
3D printed bioelectronic scaffolds support dense and viable OVCAR8 cell culture over seven days. **a)** Schematic demonstrating that cells are seeded onto bioelectronic scaffolds. Cells are seeded by applying a droplet of concentrated cell suspension to the scaffold. **b)** Representative fluorescence microscopy maximum intensity projections of Z stacks of OVCAR8 cells on scaffolds after Live/Dead™ staining (Calcein AM, green; ethidium homodimer-1, red). Thickness ∼ 124 - 140 µm. Cells proliferate, first densely covering scaffolds struts and later filling pores. **c)** Quantification of cell number over time. Dotted line represents amount of seeded cells. Mean and standard deviation presented. N=3. **d-e)** Representative scanning electron micrographs of scaffolds before (i) and seven days after cell seeding (ii). Asterisks indicate open porosity in scaffolds that becomes filled with cells.

Prior to cell seeding, scaffolds were incubated overnight in fetal bovine serum (FBS) to improve cell attachment^49,54^. High cell attachment was observed; the number of cells present after one day was nearly the same as the number seeded (105 ± 13.2%; Supplementary Figure 6). As expected, cells continuously proliferated over the seven days. In the beginning of culture, cells adhered to and proliferated on scaffold struts and by day 7, cells appeared to be densely filling the porous volume (Figure 3B). To examine this further, scanning electron microscopy revealed that the cells closed off pores and spanned multiple scaffold layers (Figure 3D-E, Supplementary Figure 7). Altogether, the cell number increased 381 ± 73.4% by day 7 (initial seeding number = 100,000 cells, Figure 3C). Overall, these results demonstrate that the bioelectronic scaffolds are suitable substrates for supporting a population of OVCAR8 cells prior to investigations of addition of chemotherapeutic compounds.

Next, before monitoring cytotoxicity after drug application, it is first necessary to determine whether the presence of cells on the scaffolds can be detected using measurements of impedance. At mid to high range frequencies of the sinusoidal voltage (10^2^ – 10^4^ Hz), cells adhered to electrodes act as insulators (high cell membrane impedance), thereby hindering ion flow and increasing the measured impedance magnitude (|Z|)^24,28^ (Figure 4A). At these frequencies, ionic current is expected to flow around the cell bodies, taking the paracellular (between cells) and subcellular (beneath cells) routes^24,28^. Therefore, changes in impedance reflect *global* changes of the cells-electrode system, including cell-cell interactions (including how tightly cells are packed together) and cell interactions with the substrate^28^. In addition to the impedance magnitude, studies of 2D impedance monitoring have elucidated that, capacitance, which is obtained from the imaginary part of the impedance, is correlated with changes in cell number at high frequencies^28,29^. At high frequencies >10^4^ Hz, the impedance of cell membrane decreases leading to current favoring capacitive coupling across the cell membrane and reducing the relative contribution of current following paracellular routes^24,28^. At these frequencies, the contribution of the capacitance of the cell membrane to the measured impedance becomes significant. Increased cell coverage of the electrode reduces the effective electrode capacitance (and thereby system capacitance) relative to a bare electrode^28^. Here, we investigate whether impedance and/or capacitance can be used to detect the presence of cells relative to a cell-free device.

**Figure 4:**
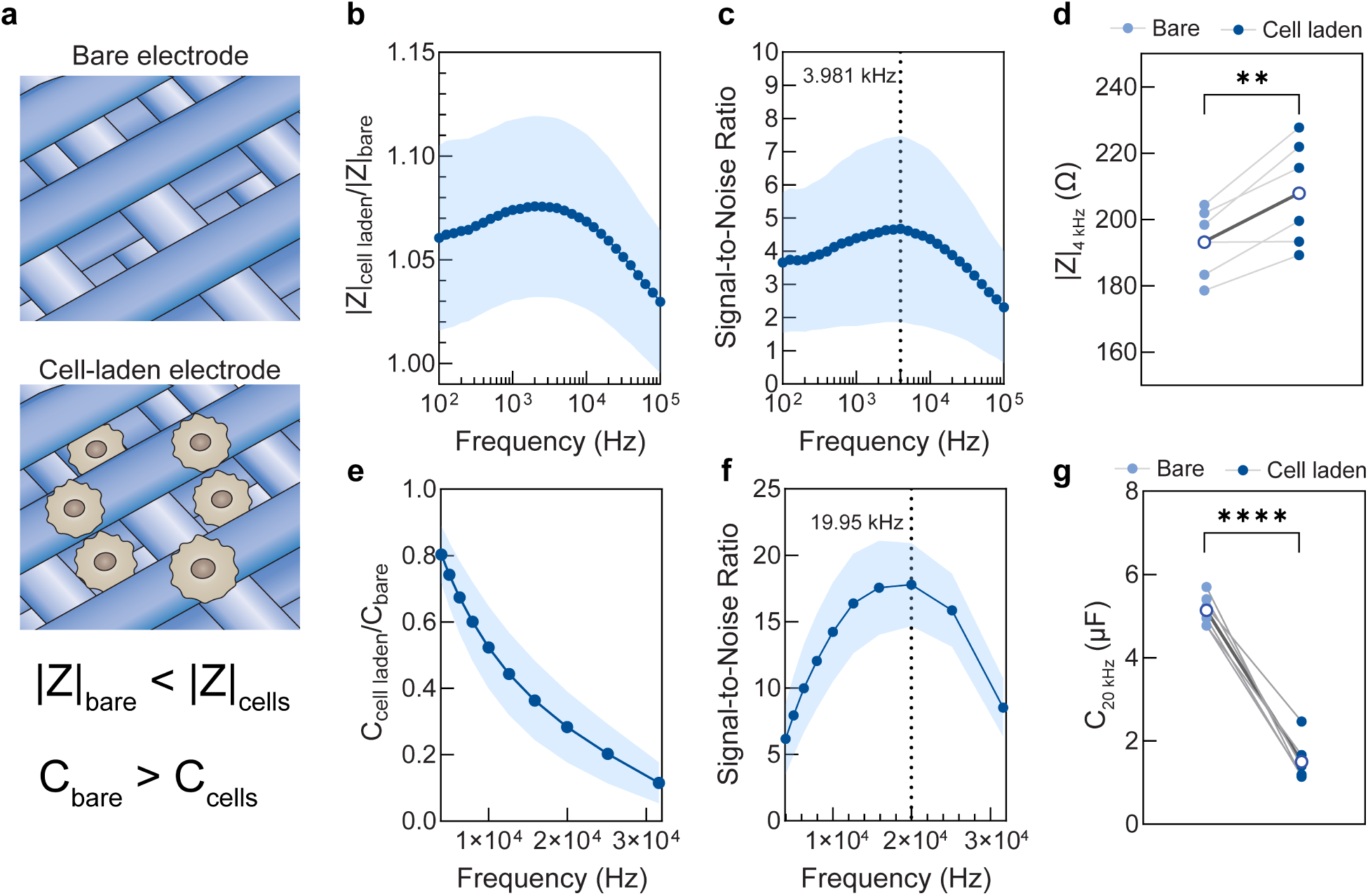
Identifying optimal frequency for the detection of cells on bioelectronic scaffold electrodes. **a)** Schematic describing the effect of cells on measured impedance and capacitance. **b)** The ratio of |Z| between cell-laden and bare scaffold electrodes at mid-range frequencies (10^2^ – 10^5^ Hz). All studied frequencies had an increased impedance with cells relative to bare electrodes (impedance ratios > 1). **c)** Signal-to-noise ratio of cell-seeded devices as a function of frequency to determine the optimal frequency for impedance analysis. The highest signal-to-noise ratio was found at 3.98 kHz (dashed line). **d)** Comparison of impedance between cell-free and cell-seeded devices at 3.98 kHz, herein labeled as 4 kHz on plots. Paired t-test performed (P = 0.0095, 95% CI [5.408, 23.77]). **e)** The ratio of C between call-laden and bare scaffold electrodes at mid-range frequencies (4 – 31.6 kHz). All examined frequencies resulted in lower capacitance with cells relative to bare electrodes (capacitance ratios < 1). **f)** Signal-to-noise ratio of cell-seeded devices as a function of frequency to determine the optimal frequency of capacitance analysis. The highest signal-to-noise ratio was found at 19.95 kHz (dashed line), herein labeled as 20 kHz on plots. **g)** Comparison of capacitance in bare and cell-laden devices at 19.95 kHz. Paired t-test performed (P < 0.0001, 95% CI [-4.306, -2.966]). Mean (line) and standard deviation (shaded area) presented for b-c, e-f (N = 6). Individual replicates presented as color points and mean presented as white circles with blue border in d & g.

Impedance was measured before and one day after seeding 250,000 cells, and capacitance was extracted from these impedance values. When examining the mid-range frequencies (∼10^2^ – 10^5^ Hz) after cell seeding, the impedance magnitude increased as expected from prior literature^28,32,45–47,53^ (Figure 4B, Supplementary Figure 8). Among this frequency range, we then sought to determine the optimal frequency for detecting cells and examined signal-to-noise ratio as a metric to do so. Although SNR analysis is a standard and widely used metric in bioanalytical instrumentation^58,59^, it is not typically applied in electrochemical impedance monitoring studies. The SNR was calculated by dividing the difference between the impedance magnitude of cell-laden and cell-free devices by the noise determined as described above (day-to-day variability, as measured by standard deviation of |Z| in cell culture conditions, Supplementary Figure 5). Typically, an SNR ≥3 is considered a good limit of detection, meaning the signals can be detected against the control, while an SNR ≥10 is considered good for the limit of quantitation, meaning the signal can be reliably quantified for precision and accuracy^60^. Performing measurements at the highest signal-to-noise ratio gives greater confidence that any observed changes in electrochemical properties can be attributed to the presence of cells rather than noise. While the SNR was similar across the frequencies, the highest SNR of 4.67 was found at 3.981 kHz (Figure 4C), and this frequency was utilized for all subsequent impedance analyses. At this frequency, the mean impedance increased by 7.49 ± 4.33% after cell seeding as compared to that of the cell-free device (Figure 4D). This increase was greater than 3X the noise defined above (2%, Supplementary Figure 5C) and thus can be attributed to the presence of cells.

Subsequently, the same analyses were performed for capacitance to identify the optimal frequency for detecting the presence of cells. When reviewing the frequency range of ≥4 kHz identified by prior literature^28,29^, the capacitance decreased with cells as expected relative to a bare electrode (Figure 4E). It should be noted that frequencies greater than 31.6 kHz were excluded from analysis (see Methods). Next, we identified the frequency that provided the highest signal-to-noise ratio in capacitance (17.8) as 19.95 kHz (Figure 4F). At this frequency, the mean capacitance decreased by 70.6 ± 9.99% (Figure 4G), a change 14 times greater than the 5% noise determined for capacitance at this frequency (Supplementary Figure 5G). In summary, these results confirm that both impedance and capacitance can be used as metrics for the detection of OVCAR8 cells on these bioelectronic scaffolds.

### Cell proliferation and carboplatin-induced cytotoxicity are monitored with bioelectronic scaffold electrodes

When a cytotoxic agent, such as a chemotherapy drug, is introduced, cell death occurs. In the context of impedance monitoring of cells, dead cells may either detach from the electrode surface or remain attached with compromised membranes. As dead cells detach, the area of bare electrode increases. If some dead cells remain attached, disintegrated cell membranes no longer effectively impede ion flow or participate in capacitive coupling^32,61^. In either case, impedance is expected to decrease and capacitance is expected to increase when cell death occurs on an electrode relative to a highly viable cell population (Figure 5A). Here, we sought to determine if the bioelectronic scaffolds could detect changes in cell number caused by cell proliferation or drug-induced cell death. The specific drug tested here was the chemotherapeutic drug carboplatin. Carboplatin is a widely used platinum-containing compound that works by damaging the DNA of cancer cells, interfering with their growth and division, ultimately leading to their death^62^. In this study, cells were cultured on bioelectronic scaffolds for 24 hours to allow for cell attachment and acclimation to the new culture environment. Carboplatin-containing media was then added to one group of the devices and was in contact with the cells for 72 hours. This timepoint is typical for evaluating drug cytotoxicity in vitro 3D cancer models since substantial cell death is expected by this time. Following this incubation, media was exchanged for drug-free media and the samples were incubated for an additional 72 hours as studies have suggested maximal drug action occurs after its removal^16,23^. A control group of devices was only subject to normal growth medium to support cell proliferation over time.

**Figure 5:**
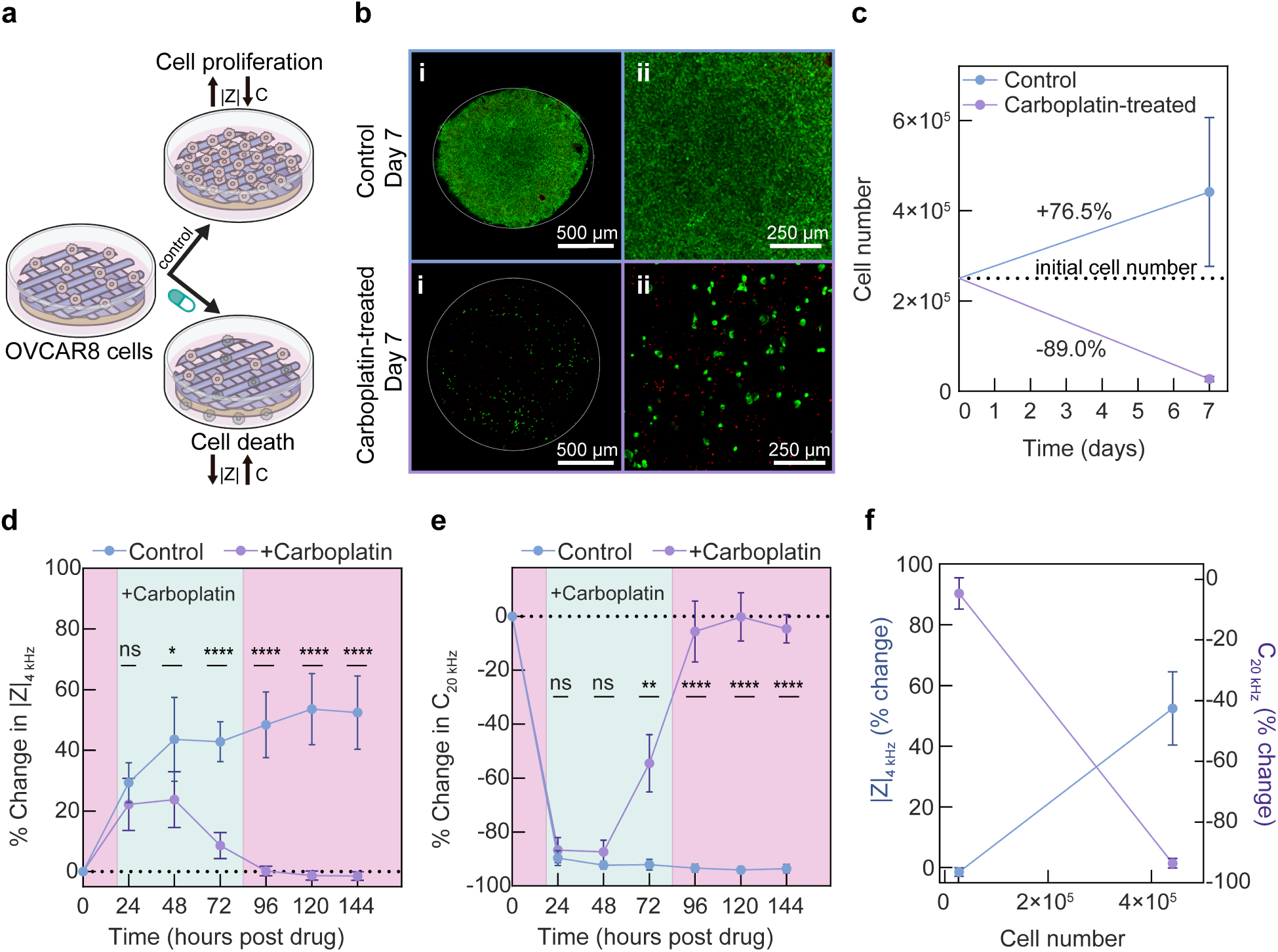
Cell proliferation and carboplatin-induced cytotoxicity are monitored with bioelectronic scaffold electrodes. **a)** Schematic describing expected impedance and capacitance changes due to cell proliferation and cytotoxicity. **b)** Representative fluorescence microscopy maximum intensity projection Z stacks of (i) tile scans and (ii) region of interests of OVCAR8 cells in the control group and carboplatin-treated group 7 days after seeding. Samples stained with Live/Dead™ staining (Calcein AM, green; ethidium homodimer-1, red). Scaffold perimeter outlined. **c)** Changes in cell number in the control and carboplatin-treated groups over 7 days. Mean and standard deviation presented, N≥3. Dotted lines represent initial number of seeded cells. **d & e)** Impedance and capacitance over time in control and drug-treated groups presented as percent change relative to their individual bare electrode values. |Z| analyzed at ∼4 kHz and C at ∼20 kHz, as determined in Figure 4. Dotted line represents the baseline bare electrode value. Mean and standard deviation presented, N≥6. Two-way analysis of variance (ANOVA) and Šidák-adjusted multiple comparison tests performed. Comparisons are done between the control and carboplatin-treated groups at each time point. ****P≤0.0001, ***P<0.001, **P≤0.01, *P≤0.05. Exact p values can be found in Supplementary Table 1. Cyan shaded region represents duration where carboplatin treatment occurred while the pink shaded area corresponds to normal growth media (RPMI 1640 with 10% FBS). **f)** Percent changes in |Z| and C relative to the value of individual bare electrodes associated with mean cell number as determined at day 7 for control and carboplatin-treated groups. Mean and standard deviation presented, N≥6.

We first confirmed that the administered dose of carboplatin caused ovarian cancer cell death on the scaffolds. In particular, we investigated 100 μM of carboplatin which falls under the reported range of IC_50_ values for platinum-resistant 3D ovarian cancer models like OVCAR8^16,63^. Here, the samples were evaluated for cell death at the end of the study (144 hours after drug treatment), with the assumption that maximal cell death would have occurred by this time point. Upon imaging with confocal microscopy and Live/Dead staining, bioelectronic scaffold electrodes were densely covered by cells in the control group, whereas in the drug-treated group, very little cell coverage was observed, confirming significant cytotoxicity caused by the drug (Figure 5B). In the drug-treated group, relatively few “red” cells were observed, suggesting that the majority of the dead cells were detached from the scaffold. Upon quantification of cell number, cell proliferation occurred as expected in the control group with a 76.5 ± 65.9% increase in cell number as compared to the seeding density (Figure 5C). In the carboplatin-treated electrodes, there was an 89.0 ± 2.56% decrease in cell number from seeding further supporting significant cell death (Figure 5C).

Impedance monitoring provides an avenue to track changes in cell behavior over time. Specifically for drug testing applications, measurements over time may provide information on the kinetics of drug effects such as when initial and maximal cell death occurs. Also, a few studies have indicated that the cytotoxic effects of carboplatin increase after drug washout^16,23^. This suggests that continuous cytotoxicity monitoring could provide insights on when this washout effect occurs. Moreover, because impedance monitoring is not a sacrificial measurement, it allows continuous measurements from the same samples, maximizing data extraction and potentially reducing the required sample size. We measured how impedance and capacitance of the bioelectronic scaffold electrodes changed over time with and without drug treatment. For impedance and capacitance, a two-way repeated measures ANOVA revealed significant main effects of time and treatment as well as a significant interaction between time and treatment (Supplementary Table 1). Pairwise comparisons were then performed with Šídák-adjusted multiple comparison test. In the control drug-free group, impedance increased over time while capacitance decreased over time (Figure 5D-E, Supplementary Figure 9). In the carboplatin-treated group, changes in impedance and capacitance between the control and drug-treated group were non-significant after 24 hours of drug incubation (Figure 5D-E, Supplementary Table 2; p = 0.5374 and p = 0.7484, for % changes in |Z| and C respectively). By 48 hours post drug addition, a significant decrease in impedance was detected, while capacitance remained non-significant when compared to the control group ( p = 0.0318 and p = 0.2128, for % changes in |Z| and C respectively). A possible explanation for these observed differences between impedance and capacitance is that the drug may have begun to influence the system in ways that are more apparent in the impedance; for example alterations in the cell-cell interactions that did not cause major cell death or electrode coverage differences. At 72 hours post-drug treatment, impedance significantly decreased by 34.2 ± 4.30% while capacitance significantly increased by 37.6 ± 10.6% compared to the control group, suggesting a substantial cytotoxic effect( p <0.0001 and p = 0.0017 for % changes in |Z| and C respectively). This result is in agreement with other studies using carboplatin where changes in cell viability are expected to be significant and thus commonly evaluated at this 72 hour timepoint^64^. After the 72 hour timepoint, the media was then exchanged to drug-free media.

During this period (96-144 hours after original drug treatment), the maximum change in impedance and capacitance was observed and |Z| and C had returned to their pre-seeding, cell-free values.

Finally, we examined the relationship between cell number with impedance or capacitance by analyzing measurements conducted at 144 hours. There was a 93.8 ± 1.45% difference in cell number between the control group and the drug treated group. For this change, impedance increased proportionally to cell number with a 54.0 ± 1.44% rise (Figure 5F). In contrast, capacitance decreased as cell number increased, with an 88.8 ± 5.22% reduction (Figure 5F). Both impedance and capacitance had a strong relationship with cell number as indicated by a Pearson’s correlation coefficient greater than ±0.81 (Supplementary Figure 10). Thus, these results show that the bioelectronic scaffolds could detect carboplatin-induced cytotoxicity in OVCAR8 cells through impedance and capacitance measurements that correspond to changes in cell number.

## Conclusions

In this work, we present bioelectronic scaffolds as devices for monitoring cancer cells by impedance. By characterizing device electrochemical properties, we determined that devices exhibit reproducible, stable impedance measurements with low noise and no apparent drift in cell culture conditions. Impedance magnitude (|Z|) and capacitance (C) were investigated as metrics for detecting cells on bioelectronic scaffolds. We utilized the signal-to-noise ratio (SNR) to select the optimal frequency for analysis which differs from prior works that majorly rely on pre-defined frequency ranges or maximum cell-free and cell-laden differences^28,65^. For capacitance in particular, the highest SNR was not found at the same frequency as the largest absolute change. At the identified frequencies, significant changes in |Z| and C indicated presence of cells. When comparing impedance and capacitance, the SNR of capacitance was more than threefold higher than that of impedance (Supplementary Figure 11). Finally, we demonstrate that the bioelectronic scaffolds can detect cytotoxicity induced by the chemotherapeutic drug carboplatin, as evidenced by a decrease in impedance and an increase in capacitance relative to a control group and confirmed by cell viability staining and cell number quantification. Measuring |Z| and C over time allowed for identification of when maximal changes occurred for both metrics when compared to drug-free controls.

Overall, this work establishes a proof-of-concept for monitoring cancer cell cytotoxicity using bioelectronic scaffolds. In the future, these scaffolds could be investigated for monitoring cytostatic effects (i.e. inhibition of proliferation), which are not readily resolved by single end-point assays^34,66^. Monitoring past typical endpoints may enable detection of drug resistance, including its onset and dose dependence, with potential relevance for clinical decision making. Beyond chemotherapeutic drugs, the scaffolds could be compatible for evaluating other treatment modalities, such as immunotherapy^35^. The non-sacrificial nature of the platform also permits complementary validation using other established assays, facilitating broader adoption of impedance-based measurements.

## Methods

### Fabrication of PEDOT:PSS hydrogel scaffolds

PEDOT:PSS hydrogel scaffolds were fabricated by adapting methods described in our previous work^49^ using a R-GEN 100 3D Bioprinter (RegenHu, Switzerland) with a mechanically driven extrusion printhead. Print designs were created using Shaper (RegenHu, Switzerland). Briefly, aqueous PEDOT:PSS dispersions (1.1-1.3 wt%, Heraeus) were vacuum filtered with 0.45 μm filters (polyethersulfone with polytetrafluoroethylene pre-filter, Environmental Express) and then mixed with the gelation agent ionic liquid (IL), 4-(3-butyl-1-imidazolio)-1-butanesulfonic acid triflate (Santa Cruz Biotechnology) at 40 mg/mL by vortex mixing followed by stirring. Inks were loaded into a Hamilton syringe customized for the R-GEN 100 printer, with a 30-gauge needle with an inner diameter 160 μm, length 25.4 mm (SAI infusion technologies). Next, ink was extruded into an agarose granular gel support consisting of 0.5 w/v% agarose granular gel mixed with 70 mg ionic liquid 4-(3-butyl-1-imidazolio)-1-butanesulfonic acid triflate (AmBeed) per 1 g agarose support medium. Cylindrical scaffolds with dimensions of 6 mm diameter, ∼1 mm thickness (6 layers), and a bounding perimeter were fabricated. Scaffolds possessed pores created by placing struts with a 150 µm center-to-center distance and an advancing angle between layers of 90°. After printing, hydrogels were incubated for 16 hours at 60 °C followed by removal of agarose by washing with deionized water^49^. To fabricate hydrogels with the desired mechanical properties, scaffolds were post-treated in 10.4 M acetic acid solution overnight, followed by de-ionized water (DI water) exchanges. In our prior work, this post processing treatment yielded a stiffness of ∼40 kPa^49^, which falls within the range known to support tumor cell culture^38,67–69^. This post-treatment resulted in material shrinkage, and therefore resulted in the following approximate final scaffold dimensions: 4 mm diameter, 600 μm thickness and 90 μm center-to-center strut pitch.

### Device fabrication

Electrode masks were designed in Adobe Illustrator and cut out of 5 mil adhesive Mylar using a CO_2_ laser cutter. Each mask included three working electrodes (3 mm diameter) and one large counter electrode (400 mm^2^ area) within the chamber slide area, connected by thin connecting lines (1 mm by 25 mm) and contact pads. For all devices, the distance between the counter electrode and working electrode was maintained as 11 mm. This Mylar mask was then carefully aligned with a clean glass slide. Gold was deposited on the masked glass slides using thermal evaporation. Chromium (10 nm) and gold (100 nm) layers were sequentially deposited using a Kurt Lesker Nano 36 Thermal Evaporator. After evaporation, the Mylar masks were removed, and the electrodes were cleaned by sequential rinsing with DI water, acetone, and IPA, followed by drying with nitrogen gas and plasma treatment for 90 seconds (Harrick Plasma PDC001) to improve surface adhesion.

To facilitate bonding of the PEDOT:PSS scaffolds to gold, an adhesion layer was made using PEDOT:PSS and 4-arm PEG thiol as follows: an adhesion layer solution – 5 w/v% of 4-arm PEG-SH (MW = 10 kDa, Laysan Bio) in PEDOT:PSS aqueous dispersion was prepared by vortex mixing for 1 minute followed by centrifugation for 30 seconds - 1 minute at 3200 xg to remove air bubbles. Next, 3 μL of adhesion layer solution were drop cast on each working electrode, followed by 10 minutes drying at 100 °C on a hot plate. After drying, the adhesion layer on the electrodes was washed in DI water with three water exchanges. After washing was completed, the samples were dried for 10 minutes at 100 °C.

Polyimide stickers (25 x 12 mm rectangles of 1 mil Kapton, silicone adhesive backing), were then used to insulate the electrode connection lines in order to define the active electrode area. Devices were next assembled into the chamber slides (Chemglass Life Sciences, 1 well). To prevent media leaking, PDMS (Krayden Dowsil Sylgard 184 elastomer kit) was applied to the silicon gaskets of chamber slides prior to assembly and allowed to cure at 60 °C for 2 hours.

To attach scaffolds to fabricated electrodes (gold + adhesion layer), 3D printed PEDOT:PSS scaffolds in water were picked up with a spatula, slightly dabbed with Kimwipes to remove excess water, and then transferred to the electrode surface. A small amount of deionized water (2 μL) was added to each scaffold to facilitate transfer. The samples were then dried for 5 minutes at 100 °C on a hot plate. To facilitate direct contact between the hot plate and the glass electrode slide, the outermost clamp of the chamber slide was removed prior to drying. After drying, the chamber slides were reassembled, and 5 mL of DI water was added to enable re-swelling of the scaffolds. The scaffolds were allowed to equilibrate overnight on an orbital shaker at 50 rpm.

### Electrochemical Characterization

Electrochemical impedance spectroscopy was performed using a potentiostat (Metrohm Autolab PGSTAT302N). As described earlier, the device setup consists of a 2-electrode setup (Figure 2A, Supplementary Figure 1). Alligator clips were used to connect the potentiostat to the device through contact pads outside the chamber well. The applied AC sinusoidal voltage was 0.01 V and all measurements were performed after an open circuit potential was established.

Electrochemical impedance spectroscopy measurements were carried out at 10 frequencies per decade. All measurements were performed with devices at room temperature (∼20-23 °C, ∼30 mins after removal from incubator held at 37 °C). For electrochemical impedance spectroscopy, a 5 mL volume of electrolyte (media or PBS at pH = 7.4) was used. Impedance magnitude (|Z|), phase, real (Z’), and imaginary (Z”) impedance components were directly extracted using Metrohm NOVA analysis software. The effective capacitance of the system was extracted from the imaginary impedance using the following equation

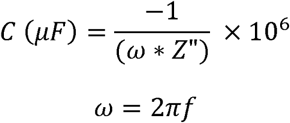

where ω is the angular frequency and f = frequency.

At high frequencies (frequencies > 31.6 kHz in our case), inductive effects from electrical wiring and connections^50,70^ become significant (Z’’>0) and the equation produces non-physical results. Therefore, frequencies greater than 31.6 kHz were excluded from capacitance analysis.

Signal-to-noise ratio *(SNR)* for impedance and capacitance was calculated by using the equations below.

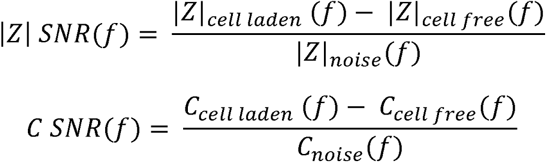

where noise is defined as the standard deviation of day-to-day variability as identified in Supplementary Figure 5.

Examination for potential drift was conducted by performing a simple linear regression of impedance or capacitance of 9 sample replicates (9 individual electrodes) over time using GraphPad Prism. Presence of drift was indicated if the slope of the regression was significantly different from 0 using an F test.

### Scanning electron microscopy (SEM)

Scaffolds were imaged using a Zeiss EVO 10 scanning electron microscope to visualize their morphologies before and after cell seeding. Scaffolds which were not bonded to gold were fixed with 2.5 v/v% glutaraldehyde and 2 v/v% paraformaldehyde in PBS for 1 hour at room temperature then overnight at 4 °C. Scaffolds were then washed with PBS, followed by serial dehydration with ethanol, and finally dehydrated using a critical point drier (K850, Electron Microscopy Sciences). Scaffolds bonded to gold were first gently scraped using scalpels to detach from gold, followed by complete dehydration through lyophilization. To perform SEM imaging, the dried samples were mounted to stubs using carbon tape followed by sputtering with 5 nm gold. All SEM images were captured at 20 kV with a working distance ranging from 9 to 40 mm.

### Scaffold and device preparation for in vitro studies

Prior to in vitro studies, scaffolds were transferred to a 12-well plate, disinfected in 70% ethanol overnight in a biosafety cabinet, followed by UV exposure for 30 minutes-1 hour and then washed in sterile 1X phosphate-buffered saline (PBS) with 1% penicillin–streptomycin (Pen/Strep; 10,000 U mL^-1^ penicillin and 10,000 μg/mL streptomycin) for 15 minutes. Afterwards, scaffolds were transferred to sterile 12-well plate coated with 2 wt% agarose which provides a non-cell adhesive surface beneath the scaffolds followed by overnight incubation in PBS+Pen/Strep. For devices, the inside of the chamber slides was disinfected similarly to scaffolds. Chamber slide lids were also submerged in 70% ethanol overnight and then allowed to dry in the biosafety cabinet. Next, scaffolds and devices were prepared for cell seeding by submerging in fetal bovine serum (FBS) overnight in the incubator (at 37 °C, 95% relative humidity, and 5% CO_2_). After incubation in FBS, samples were washed once with PBS+Pen/Step for 15 minutes to remove residual FBS. Finally, samples were submerged in cell culture media (RPMI 1640 reconstituted with 10% FBS and 1% Pen/Step) and placed in incubator until cell seeding or impedance monitoring experiments (∼1-3 days). All samples were handled aseptically in a biosafety cabinet during preparation and for the duration of the study.

### Cell seeding

Human immortalized ovarian cancer cell line OVCAR8was cultured, expanded on T75 flasks and used at passage 36-38. Immediately prior to cell seeding scaffolds and devices (lid open) were dried for 15 and 10 minutes respectively in the biosafety cabinet to allow scaffolds to later wick up the applied cell suspension. 100,000 cells were seeded onto each scaffold by depositing 3 µL of prepared cell suspension on dried scaffolds (Figure 3). For the scaffold working electrodes, 250,000 cells were similarly seeded using 3 µL suspension volume per scaffold (Figures 4 and 5). After seeding, samples were transferred to an incubator for 45 minutes to allow for cell attachment. Cell culture media was then added to each well or chamber (4 mL and 5 mL for scaffolds in well plates or assembled devices, respectively). Cell culture media was changed every other day.

### Cytotoxicity of Applied Chemotherapy Drug

Cells were seeded on the scaffold working electrodes at a density of 83.33 million cells/mL (250,000 cells per scaffold) and allowed to attach and proliferate for 24 hours undisturbed. 10 mM carboplatin stock solution was prepared in de-ionized water using manufacturer’s instructions (Selleckchem). Drug-free cell culture media was then replaced with 100 μM of carboplatin solution in media prepared from stock solution. After 72 hours of incubation, drug-containing media was replaced with drug-free media and studies continued for an additional 72 hours. Control samples were maintained in drug-free media and media was exchanged after 24 hours, and then after 96 hours to match the media replacement in treatment condition.

Impedance measurements were performed every 24 hours on devices. At the end of the studies, end point analysis of cell imaging and cell number quantification were performed on devices, described as follows.

### Cell analysis

Cell-seeded scaffolds were evaluated for cell attachment through staining (LIVE/DEAD™ kit (Invitrogen, 2 µM Calcein AM, 8 µM of ethidium homodimer-1) and fluorescence microscopy. Each scaffold was transferred to a confocal glass bottom dish and stained for at least 30 minutes. Imaging was performed with a Leica SP8 Lightning confocal microscope. For devices, the scaffold electrodes were similarly stained in the chamber well. After staining for at least 30 minutes, staining solution was removed, and chamber slide was disassembled resulting in just glass slides with scaffolds attached. These glass slides were then inverted on well plates with rectangular wells equipped with 450 μm thick PDMS film to help suspend the glass and prevent scaffolds from being crushed by the weight of the glass.

To measure cell number, the Quant-iT™ PicoGreen™ (Invitrogen) assay was used. Cell-seeded scaffolds were collected in microtubes and frozen at -20 °C at select time points until time of the assay. Scaffolds adhered to electrodes were carefully scraped off the glass substrate using a scalpel and similarly collected in microtubes. Aliquots of the cell seeding suspension were also collected. All samples were then lyophilized and digested for DNA extraction in Proteinase K solution overnight at 60 °C. Afterwards, the Picogreen™ assay was then run to quantify DNA content on scaffolds using the manufacturer’s instructions. Cell attachment efficiency (%) was determined by dividing the cells present after one day by the cell number from the cell seeding suspension and multiplying by 100.

### Statistical analysis

For statistical analysis, t tests, F test, ordinary and repeated measures one-way ANOVA with Šidák’s-adjusted multiple comparison tests and repeated measures two-way ANOVA with Šidák’s-adjusted multiple comparison tests were performed with GraphPad Prism. Results are presented as means with positive and negative standard deviation unless otherwise noted.

Differences were categorized as statistically significant for p < 0.05. Pearson correlation test was performed using GraphPad Prism.

## Data availability

The data that support the findings of this study are available from the corresponding author upon reasonable request.

## Author Contributions

S.S.O., M.M.M. and A.L.R. envisioned this study and wrote this manuscript. S.S.O., S.K.M., J.P., T.L., E.L., I.S., M.M.M., and A.L.R. were involved in data interpretation and statistical analysis.

S.S.O, S.K.M and M.S. were involved in device development. S.S.O. conducted all cell culture experiments, confocal imaging, impedance data collection and analysis for cell-based impedance experiments. S.S.O. and S.K.M. performed device characterization. T.L. & S.S.O. performed SEM. S.S.O., S.K.M. and J.P. designed schematics. A.S. was involved in methods development for OVCAR8 cell culture and chemotherapy studies. A.S. & A.P.G. were involved in technical discussions. S.S.O., S.K.M., J.P., T.L., J.S.Y., C.P.O., A.P.G., C.J.V.E. and Y.W. contributed to material and device fabrication.

## Acknowledgements

This research was supported in part by Washington University in St. Louis through the Women’s Health Technologies Collaboration Initiation Grant and the Ovarian Cancer Research Innovation Fund Grant. The authors further acknowledge support from the National Science Foundation through CAREER GR #0039349 and FR #2319060. S.S.O. acknowledges support from the McDonnell International Scholars Academy. M.M.M reports funding from the Damon Runyon Research Foundation, the Reproductive Scientist Development Program (RSDP) supported by the Gynecologic Oncology Group Foundation, the NCI Early-stage surgeon scientist program, the Victoria Secret Global Fund for Women’s Cancers Career Development Award in partnership with Pelotonia and AACR, the Pilot Translational and Clinical Studies function of the Washington University Institute of Clinical and Translational Sciences, and the Foundation for Barnes-Jewish Hospital, the Foundation for Women’s Cancer, and the American Cancer Society. M.M.M. reports personal fees from Valinor Discovery and Bioscent Diagnostics. The authors would like to thank Dianne Duncan (Director of the Washington University in St. Louis Biology Department Imaging Facility) for technical assistance regarding imaging. The authors also acknowledge assistance from staff and use of instruments from the Institute of Materials Science and Engineering, Department of Mechanical Engineering and Materials Science at Washington University in St. Louis. The authors also acknowledge Prof. Nathaniel Huebsch for providing access to the plasma cleaner & lyophilizer, and Prof. Cory Berkland for providing access to the plate reader. The authors acknowledge the use of AI tools ChatGPT and Claude (2026) to improve grammatic clarity, conciseness, flow and accuracy in the Abstract, Result and Discussion sections. All AI-assisted text was reviewed and revised by the authors to ensure accuracy and clarity of meaning.

## Ethics declaration Competing interests

S.S.O., M.M.M. and A.L.R. are inventors of a U.S. patent application that covers the use of the bioelectronic scaffolds for drug monitoring. All other authors declare no competing interests.

## References

1. Bray, F., et al. Global cancer statistics 2022: GLOBOCAN estimates of incidence and mortality worldwide for 36 cancers in 185 countries. CA: A Cancer Journal for Clinicians 74, 229–263 (2024).

2. Bhat, G. R. et al. Cancer cell plasticity: from cellular, molecular, and genetic mechanisms to tumor heterogeneity and drug resistance. Cancer Metastasis Rev 43, 197–228 (2024).

3. LeSavage, B. L., Suhar, R. A., Broguiere, N., Lutolf, M. P. & Heilshorn, S. C. Next-generation cancer organoids. Nat. Mater. 21, 143–159 (2022).

4. Wensink, G. E. et al. Patient-derived organoids as a predictive biomarker for treatment response in cancer patients. *npj Precis*. Onc. 5, 1–13 (2021).

5. Thorel, L. et al. Patient-derived tumor organoids: a new avenue for preclinical research and precision medicine in oncology. Exp Mol Med 56, 1531–1551 (2024).

6. Kim, J., Koo, B.-K. & Knoblich, J. A. Human organoids: model systems for human biology and medicine. Nat Rev Mol Cell Biol 21, 571–584 (2020).

7. Kim, J., Koo, B.-K. & Knoblich, J. A. Human organoids: model systems for human biology and medicine. Nat Rev Mol Cell Biol 21, 571–584 (2020).

8. Griffith, L. G. & Swartz, M. A. Capturing complex 3D tissue physiology in vitro. Nat Rev Mol Cell Biol 7, 211–224 (2006).

9. Moss, S. P., Bakirci, E. & Feinberg, A. W. Engineering the 3D structure of organoids. Stem Cell Reports 20, 102379 (2025).

10. Nwokoye, P. N. & Abilez, O. J. Bioengineering methods for vascularizing organoids. Cell Reports Methods 4, 100779 (2024).

11. Zhao, Y. et al. Integrating organoids and organ-on-a-chip devices. Nat Rev Bioeng 2, 588–608 (2024).

12. Abuwatfa, W. H., Pitt, W. G. & Husseini, G. A. Scaffold-based 3D cell culture models in cancer research. Journal of Biomedical Science 31, 7 (2024).

13. Tong, L. et al. Patient-derived organoids in precision cancer medicine. Med 5, 1351–1377 (2024).

14. Foo, M. A. et al. Clinical translation of patient-derived tumour organoids- bottlenecks and strategies. Biomarker Research 10, 10 (2022).

15. Qu, S. et al. Patient-derived organoids in human cancer: a platform for fundamental research and precision medicine. Mol Biomed 5, 6 (2024).

16. Brodeur, M. N. et al. Carboplatin response in preclinical models for ovarian cancer: comparison of 2D monolayers, spheroids, ex vivo tumors and in vivo models. Sci Rep 11, 18183 (2021).

17. Nunes, A. S., Barros, A. S., Costa, E. C., Moreira, A. F. & Correia, I. J. 3D tumor spheroids as in vitro models to mimic in vivo human solid tumors resistance to therapeutic drugs. Biotechnology and Bioengineering 116, 206–226 (2019).

18. Astashkina, A., Mann, B. & Grainger, D. W. A critical evaluation of in vitro cell culture models for high-throughput drug screening and toxicity. Pharmacology & Therapeutics 134, 82–106 (2012).

19. Lynch, C. et al. High-Throughput Screening to Advance In Vitro Toxicology: Accomplishments, Challenges, and Future Directions. Annual Review of Pharmacology and Toxicology 64, 191–209 (2024).

20. Sazonova, E. V., Chesnokov, M. S., Zhivotovsky, B. & Kopeina, G. S. Drug toxicity assessment: cell proliferation versus cell death. Cell Death Discov. 8, 417 (2022).

21. Cortesi, M. & Giordano, E. Non-destructive monitoring of 3D cell cultures: new technologies and applications. PEERJ 10, (2022).

22. Pérez-Velázquez, J. & Rejniak, K. A. Drug-Induced Resistance in Micrometastases: Analysis of Spatio-Temporal Cell Lineages. Front. Physiol. 11, (2020).

23. Jabs, J. et al. Screening drug effects in patient-derived cancer cells links organoid responses to genome alterations. Molecular Systems Biology 13, 955 (2017).

24. Moghtaderi, H. et al. Electric cell-substrate impedance sensing in cancer research: An in-depth exploration of impedance sensing for profiling cancer cell behavior. Sensors and Actuators Reports 7, 100188 (2024).

25. Opp, D. et al. Use of electric cell-substrate impedance sensing to assess in vitro cytotoxicity. Biosens Bioelectron 24, 2625–2629 (2009).

26. Xing, J. Z. et al. Dynamic Monitoring of Cytotoxicity on Microelectronic Sensors. Chem. Res. Toxicol. 18, 154–161 (2005).

27. Ngoc Le, H. T., Kim, J., Park, J. & Cho, S. A Review of Electrical Impedance Characterization of Cells for Label-Free and Real-Time Assays. BioChip J 13, 295–305 (2019).

28. Stolwijk, J. A., Matrougui, K., Renken, C. W. & Trebak, M. Impedance analysis of GPCR-mediated changes in endothelial barrier function: overview, and fundamental considerations for stable and reproducible measurements. Pflugers Arch 467, 2193–2218 (2015).

29. Wegener, J., Keese, C. R. & Giaever, I. Electric Cell–Substrate Impedance Sensing (ECIS) as a Noninvasive Means to Monitor the Kinetics of Cell Spreading to Artificial Surfaces. Experimental Cell Research 259, 158–166 (2000).

30. Xu, Y. et al. A review of impedance measurements of whole cells. Biosensors and Bioelectronics 77, 824–836 (2016).

31. Abasi, S., Aggas, J. R., Garayar-Leyva, G. G., Walther, B. K. & Guiseppi-Elie, A. Bioelectrical Impedance Spectroscopy for Monitoring Mammalian Cells and Tissues under Different Frequency Domains: A Review. ACS Meas. Sci. Au 2, 495–516 (2022).

32. Anh-Nguyen, T., Tiberius, B., Pliquett, U. & Urban, G. A. An impedance biosensor for monitoring cancer cell attachment, spreading and drug-induced apoptosis. Sensors and Actuators A: Physical 241, 231–237 (2016).

33. Pradhan, R., Mandal, M., Mitra, A. & Das, S. Monitoring cellular activities of cancer cells using impedance sensing devices. Sensors and Actuators B: Chemical 193, 478–483 (2014).

34. Limame, R. et al. Comparative Analysis of Dynamic Cell Viability, Migration and Invasion Assessments by Novel Real-Time Technology and Classic Endpoint Assays. PLOS ONE 7, e46536 (2012).

35. Kiesgen, S., Messinger, J. C., Chintala, N. K., Tano, Z. & Adusumilli, P. S. Comparative analysis of assays to measure CAR T-cell-mediated cytotoxicity. Nat Protoc 16, 1331–1342 (2021).

36. De León, S., Pupovac, A. & McArthur, S. Three-Dimensional (3D) cell culture monitoring: Opportunities and challenges for impedance spectroscopy. BIOTECHNOLOGY AND BIOENGINEERING 117, 1230–1240 (2020).

37. Caliari, S. R. & Burdick, J. A. A practical guide to hydrogels for cell culture. Nat Methods 13, 405–414 (2016).

38. Zhang, M. & Zhang, B. Extracellular matrix stiffness: mechanisms in tumor progression and therapeutic potential in cancer. Experimental Hematology & Oncology 14, 54 (2025).

39. Pan, Y. et al. 3D microgroove electrical impedance sensing to examine 3D cell cultures for antineoplastic drug assessment. Microsyst Nanoeng 6, 1–10 (2020).

40. Lei, K. F., Wu, M.-H., Hsu, C.-W. & Chen, Y.-D. Electrical Impedance Determination of Cancer Cell Viability in a 3-Dimensional Cell Culture Microfluidic Chip. International Journal of Electrochemical Science 7, 12817–12828 (2012).

41. Lei, K. F., Wu, M.-H., Hsu, C.-W. & Lin, C.-Y. Quantification of Cell Number in 3-Dimensional Cell Culture Construct by Impedance Measurement using Microfluidic Technology. International Journal of Electrochemical Science 7, 8848–8858 (2012).

42. Pan, Y. et al. 3D cell-based biosensor for cell viability and drug assessment by 3D electric cell/matrigel-substrate impedance sensing. Biosensors and Bioelectronics 130, 344–351 (2019).

43. Qiu, Y. et al. Vertical impedance electrode array for spatiotemporal dynamics monitoring of 3D cells under drug diffusion effect. ISCIENCE 26, (2023).

44. Lei, K. F., Wu, M.-H., Hsu, C.-W. & Chen, Y.-D. Real-time and non-invasive impedimetric monitoring of cell proliferation and chemosensitivity in a perfusion 3D cell culture microfluidic chip. Biosensors and Bioelectronics 51, 16–21 (2014).

45. Savva, A. et al. 3D organic bioelectronics for electrical monitoring of human adult stem cells. Mater. Horiz. 10, 3589–3600 (2023).

46. Pitsalidis, C. et al. Organic electronic transmembrane device for hosting and monitoring 3D cell cultures. Sci Adv 8, eabo4761 (2022).

47. Inal, S. et al. Conducting Polymer Scaffolds for Hosting and Monitoring 3D Cell Culture. Advanced Biosystems 1, 1700052 (2017).

48. Moysidou, C.-M. et al. 3D Bioelectronic Model of the Human Intestine. Advanced Biology 5, 2000306 (2021).

49. Okafor, S. S. et al. 3D Printed Bioelectronic Scaffolds with Soft Tissue-Like Stiffness. Advanced Materials Technologies 10, 2401528 (2025).

50. Lazanas, A. Ch. & Prodromidis, M. I. Electrochemical Impedance Spectroscopy─A Tutorial. ACS Meas. Sci. Au 3, 162–193 (2023).

51. Bailleul, A. et al. In vitro impedance spectroscopy: A MEA-based measurement bench for myoblasts cultures monitoring. in 2021 XXXVI Conference on Design of Circuits and Integrated Systems (DCIS) 1–6 (2021). doi:10.1109/DCIS53048.2021.9666172.

52. ASSOCIATION OF PUBLIC HEALTH LABORATORIES. Verification and Validation Toolkit: Determining Performance Characteristics of Quantitative Assays. 1–6 http://www.aphl.org (2024).

53. Arndt, S., Seebach, J., Psathaki, K., Galla, H.-J. & Wegener, J. Bioelectrical impedance assay to monitor changes in cell shape during apoptosis. Biosensors and Bioelectronics 19, 583–594 (2004).

54. Goestenkors, A. P. et al. Manipulation of cross-linking in PEDOT:PSS hydrogels for biointerfacing. J. Mater. Chem. B 11, 11357–11371 (2023).

55. Newbold, C. et al. Changes in biphasic electrode impedance with protein adsorption and cell growth. J. Neural Eng. 7, 056011 (2010).

56. Schab, A. et al. Replication stress marker phospho-RPA2 predicts response to platinum and PARP inhibitors in homologous recombination-proficient ovarian cancer. 2024.11.21.624682 Preprint at 10.1101/2024.11.21.624682 (2024).

57. Hallas-Potts, A., Dawson, J. C. & Herrington, C. S. Ovarian cancer cell lines derived from non-serous carcinomas migrate and invade more aggressively than those derived from high-grade serous carcinomas. Sci Rep 9, 5515 (2019).

58. Li, C. et al. Towards Higher Sensitivity of Mass Spectrometry: A Perspective From the Mass Analyzers. Front Chem 9, 813359 (2021).

59. Suarez-Perez, A. et al. Quantification of Signal-to-Noise Ratio in Cerebral Cortex Recordings Using Flexible MEAs With Co-localized Platinum Black, Carbon Nanotubes, and Gold Electrodes. Front Neurosci 12, 862 (2018).

60. International Council for Harmonisation of Technical Requirements for Pharmaceuticals for Human Use. Validation of Analytical Procedures Q2(R2). https://www.ich.org/page/quality-guidelines (2023).

61. Lei, K. F., Liu, T.-K. & Tsang, N.-M. Towards a high throughput impedimetric screening of chemosensitivity of cancer cells suspended in hydrogel and cultured in a paper substrate. Biosensors and Bioelectronics 100, 355–360 (2018).

62. Ho, G. Y., Woodward, N. & Coward, J. I. G. Cisplatin versus carboplatin: comparative review of therapeutic management in solid malignancies. Critical Reviews in Oncology/Hematology 102, 37–46 (2016).

63. Singh, T. et al. Efficacy of birinapant in combination with carboplatin in targeting platinum-resistant epithelial ovarian cancers. Int J Oncol 60, 35 (2022).

64. Cavarzerani, E., Caligiuri, I., Bartoletti, M., Canzonieri, V. & Rizzolio, F. 3D dynamic cultures of HGSOC organoids to model innovative and standard therapies. Front. Bioeng. Biotechnol. 11, (2023).

65. Ebrahim, A. S. et al. Functional optimization of electric cell-substrate impedance sensing (ECIS) using human corneal epithelial cells. Sci Rep 12, 14126 (2022).

66. Single, A., Beetham, H., Telford, B. J., Guilford, P. & Chen, A. A Comparison of Real-Time and Endpoint Cell Viability Assays for Improved Synthetic Lethal Drug Validation. SLAS Discovery 20, 1286–1293 (2015).

67. Sievers, J., Mahajan, V., Welzel, P. B., Werner, C. & Taubenberger, A. Precision Hydrogels for the Study of Cancer Cell Mechanobiology. Advanced Healthcare Materials 12, 2202514 (2023).

68. Wei, X. et al. TAGLN mediated stiffness-regulated ovarian cancer progression via RhoA/ROCK pathway. Journal of Experimental & Clinical Cancer Research 40, 292 (2021).

69. Jiang, Y. et al. Targeting extracellular matrix stiffness and mechanotransducers to improve cancer therapy. Journal of Hematology & Oncology 15, 34 (2022).

70. Sosa Gallardo, A. F. & Provis, J. L. Electrochemical cell design and impedance spectroscopy of cement hydration. J Mater Sci 56, 1203–1220 (2021).

