## Supplementary Figure for "3D Printed Bioelectronic Scaffolds for Impedance-based Cytotoxicity Monitoring of In Vitro Cancer Models"

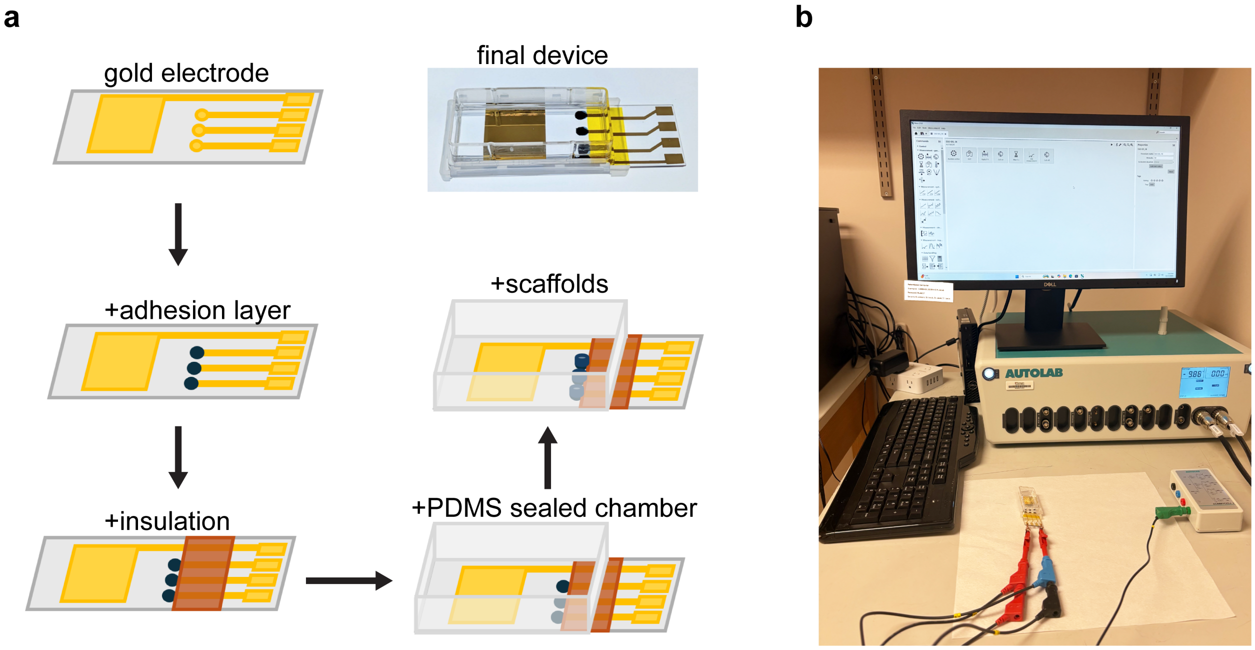


**Supplementary Figure 1. Fabrication and characterization of bioelectronic scaffold devices. a)** Schematic of fabrication process of devices. **b)** Image of device connected to instrumentation (potentiostat) for measurements.

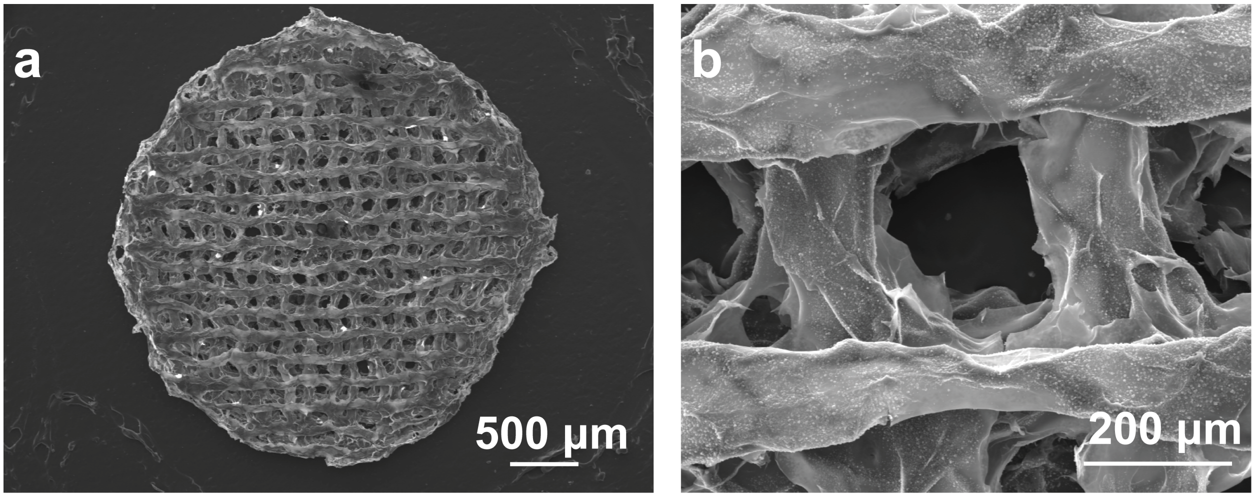


**Supplementary Figure 2. a-b)** Scanning electron micrographs of scaffolds after adhesion to gold at different magnifications. 3D printed microporous structure is maintained after adhesion to substrate.

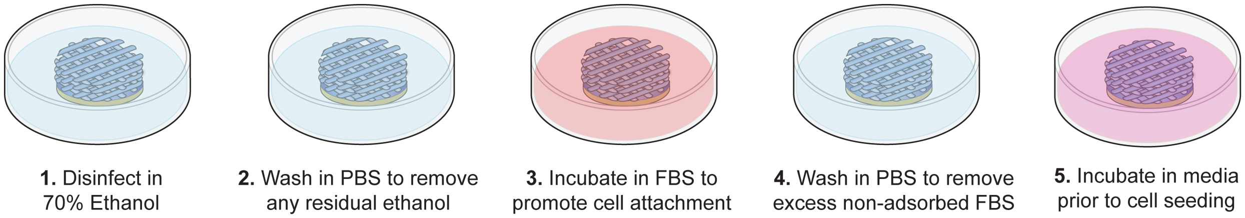


**Supplementary Figure 3. Preparative processes for cell culture studies.** PBS = phosphate-buffered saline solution. FBS = fetal bovine serum.

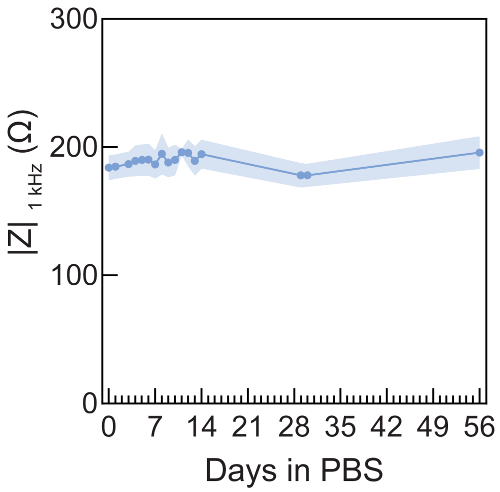


**Supplementary Figure 4. Characterization of scaffold electrode impedance in phosphate-buffered saline at room temperature over time.** Mean and standard deviation presented. N=3. A one-way ANOVA determined that the impedance magnitude did not significantly change over 56 days ($F = 0.7631$, $p = 0.7128$).

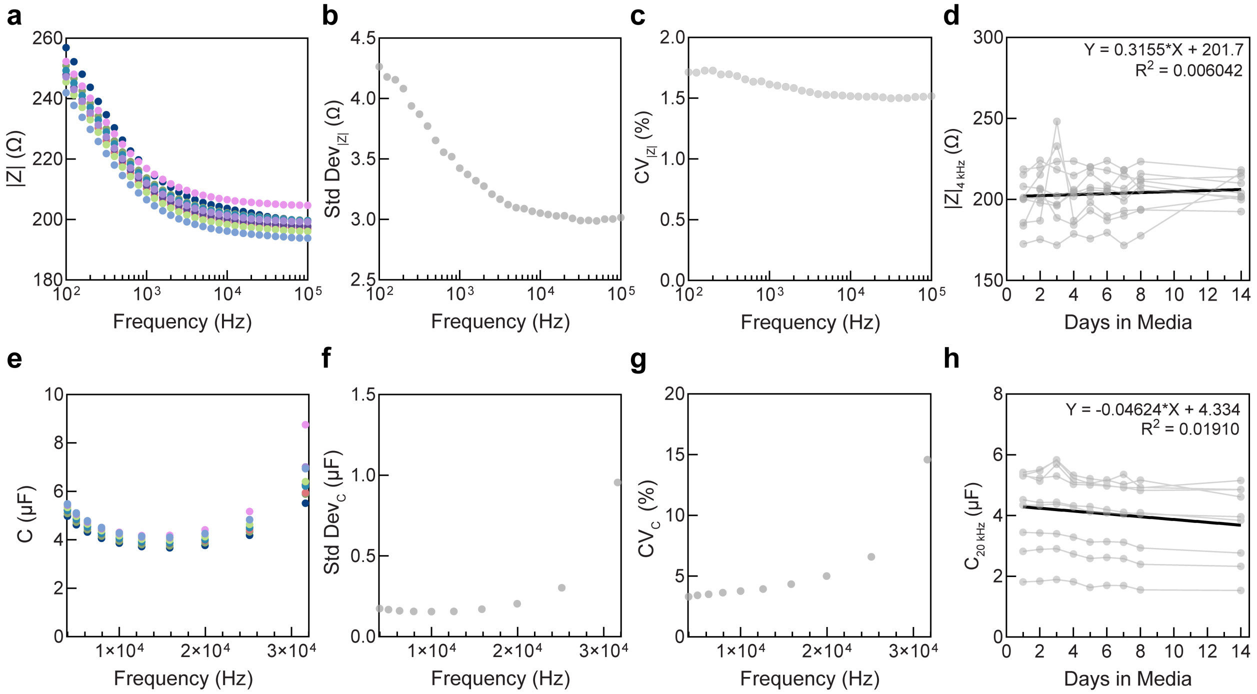


**Supplementary Figure 5: Scaffold electrode impedance and capacitance over time in cell culture conditions for characterization of noise and drift. a)** Mean impedance of electrodes over time at different frequencies. Color indicates different days in media. **b)** Standard deviation of impedance presented in (a). **c)** Coefficient of variation of impedance presented in (a). **d)** Linear regression analysis (black line) of individual electrode impedance (grey) at ~4 kHz (optimal frequency as identified in Figure 4). Fit parameters presented in the plot. An F-test comparing the fitted slope (0.3155) to a slope of 0 indicated no statistically significant evidence of drift (F = 0.4802, p=0.4903). **e)** Mean capacitance of electrodes over time at different frequencies. Color indicates different days in media. **f)** Standard deviation of capacitance presented in (e). **g)** Coefficient of variation of capacitance presented in (e). **h)** Linear regression analysis (black line) of individual electrode capacitance (grey) at ~20 kHz (optimal frequency as identified in Figure 4). Fit parameters presented in the plot. An F-test comparing the fitted slope (-0.04624) to a slope of 0 indicated no statistically significant evidence of drift (F = 1.539, p=0.2185). N=9.

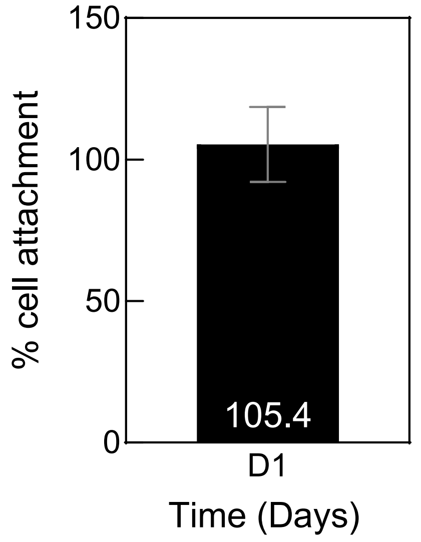


**Supplementary Figure 6: Quantification of OVCAR8 cell number on bioelectronic scaffolds one day after cell seeding.** Cells quantified as a percentage of the initial seed cell number. Mean and standard deviation presented. N=3.

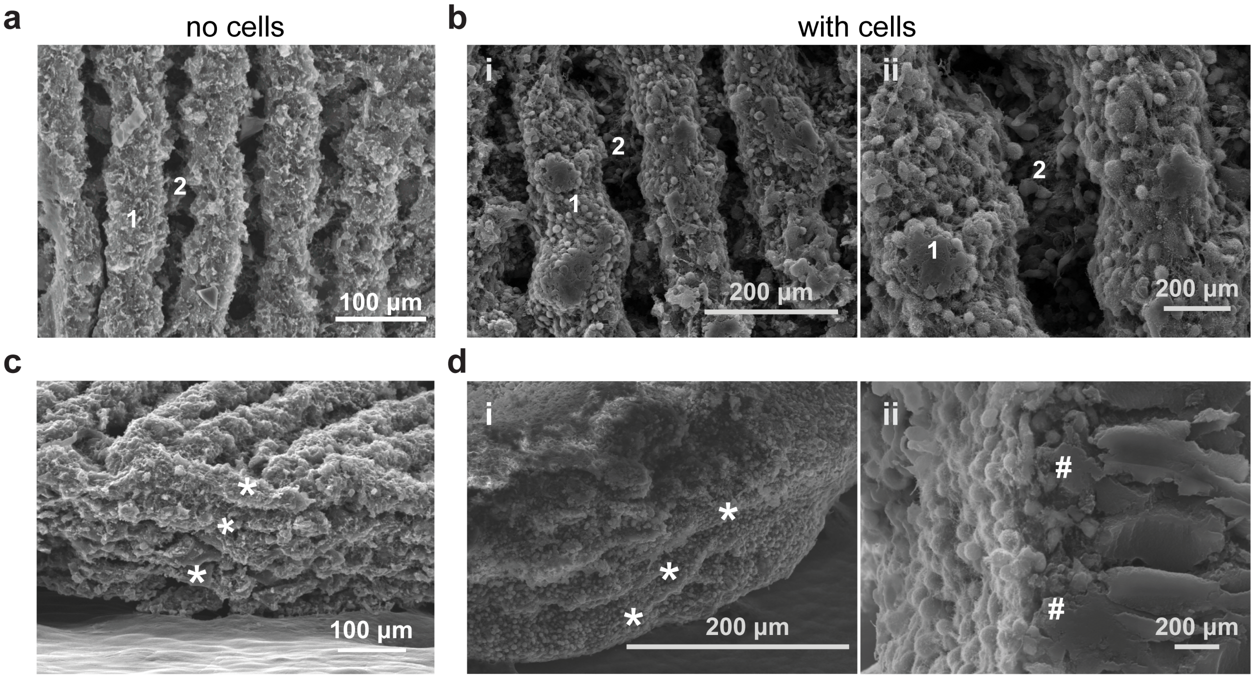


**Supplementary Figure 7: Cells proliferate into pores and across layers of scaffolds.** Top view scanning electron micrographs of scaffold **(a)** before cell seeding and **(b)** seven days after seeding with OVCAR8 cells. Visible printed layers identified as numbers 1 and 2. The interconnected porosity of the scaffolds enable the OVCAR8 cells to enter the scaffold upon seeding and attach throughout the scaffold volume. Side view scanning electron micrographs of scaffolds before cell seeding **(c)** and seven days after seeding with OVCAR8 cells **(d)**. Printed scaffold struts become difficult to discern as cell densely fill and surround the scaffold. Cells are uniformly observed across several scaffold layers (asterisks indicate discernable scaffold layers). OVCAR8 cells are observed as multiple cell layers on scaffold struts (# indicate the scaffold material).

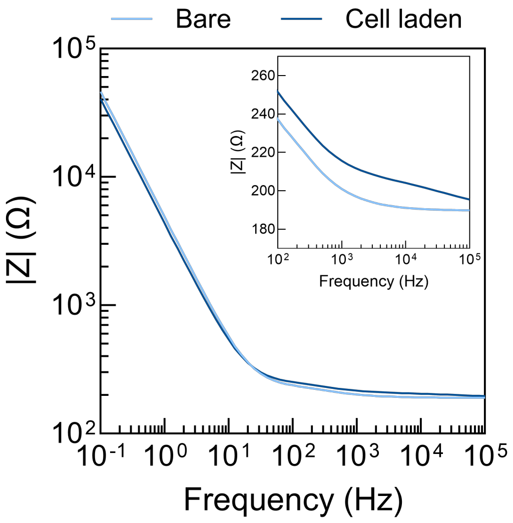


**Supplementary Figure 8: Representative Bode |Z| plot of bare and cell-laden scaffolds one day after seeding 250,000 OVCAR8 cells.** Inset shows plot zoomed in on $10^2 – 10^5$ frequencies.

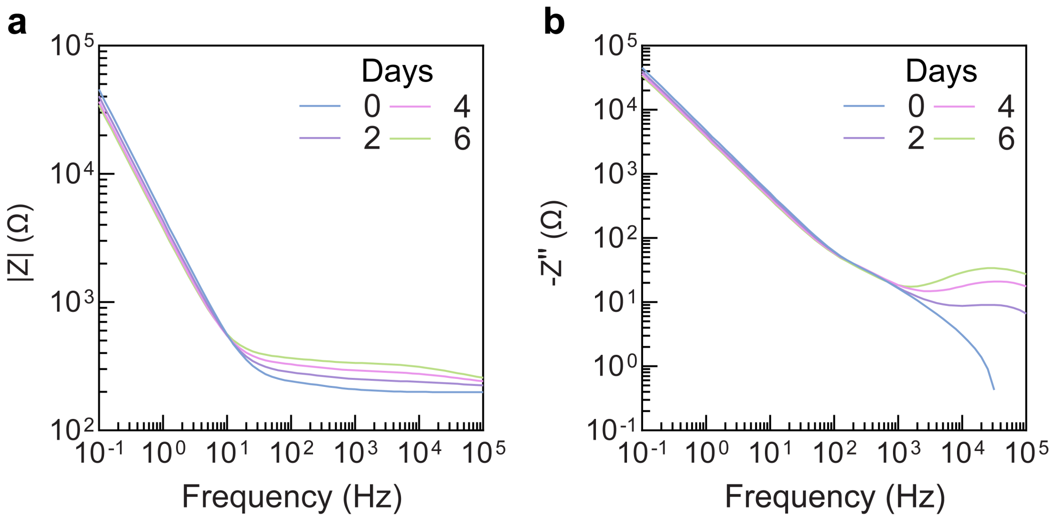


**Supplementary Figure 9: Representative Bode plots of one electrode before seeding and over time after OVCAR8 cell seeding. (a)** Impedance magnitude (|Z|) plot over time. |Z| increased in the mid frequency range ($10^2 – 10^5$ Hz) with cell proliferation. **(b)** Plot of the imaginary component of impedance over time. Z'' increased in the high frequency regions >4 kHz with cell proliferation. Capacitance is obtained from Z" using the equation in the Methods section.

**Supplementary Table 1: Results of two-way analysis of variance (ANOVA) examining the effects of time and drug treatment on impedance and capacitance.**

| Effect | \|Z\| (F value, P value) | C (F value, P value) |
|---|---|---|
| Time and treatment interaction | F = 53.61, P<0.0001 | F = 649.4, P<0.0001 |
| Time effect | F = 6.411, P = 0.0068 | F = 570.6, P<0.0001 |
| Treatment effect | F = 79.27, P<0.0001 | F = 534.6, P<0.0001 |

**Supplementary Table 2: P values and 95% confidence interval values from Šidák-adjusted multiple comparisons test following two-way ANOVA in Supplementary Table 1.** Comparisons are done between control and carboplatin-treated groups at each time point for changes in impedance and capacitance.

| Time (h) | \|Z\| (P Value, 95% CI) | C (P Value, 95% CI) |
|---|---|---|
| 24 | P = 0.5931, 95% CI [-6.847, 21.13] | P = 0.8001, 95% CI [-10.40, 4.539] |
| 48 | P = 0.0369, 95% CI [1.430, 38.32] | P = 0.2435, 95% CI [-12.02, 2.283] |
| 72 | P < 0.0001, 95% CI [25.48, 42.92] | P = 0.0020, 95% CI [-55.56. -19.54] |
| 96 | P < 0.0001, 95% CI [35.70, 60.72] | P < 0.0001, 95% CI [-107.0, -68.41] |
| 120 | P < 0.0001, 95% CI [41.35, 68.30] | P < 0.0001, 95% CI [-109.1, -78.44] |
| 144 | P < 0.0001, 95% CI [40.05, 67.87] | P < 0.0001, 95% CI [-97.51, -80.13] |

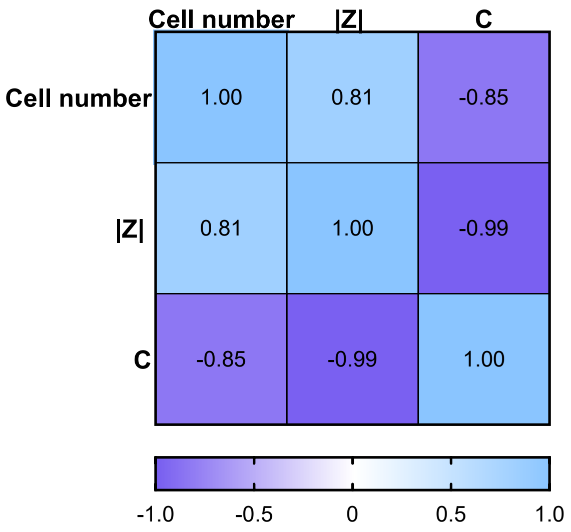


**Supplementary Figure 10: Correlation matrix comparing changes in cell number, impedance, and capacitance at the end of the study, 144 hours after drug application.**

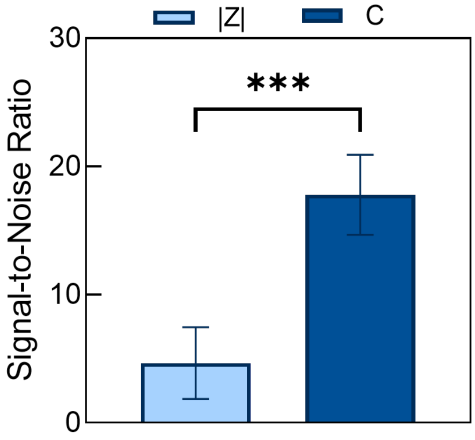


**Supplementary Figure 11: Comparison of signal to noise ratio of impedance and capacitance one day post seeding.** Mean and standard deviation presented. N=6. Paired t test, P = 0.0002, 95% CI [9.615, 16.62].
